# Reconciling multiple views of global marine plankton diversity

**DOI:** 10.64898/2026.11.25.767842

**Authors:** Alexandre Schickele, Corentin Clerc, Fabio Benedetti, Virginie Sonnet, Meike Vogt

## Abstract

Marine plankton diversity influences key ecosystem properties, including primary production and carbon export, and is monitored using diverse observation systems. Yet, it remains unclear whether global diversity patterns from different observation streams converge, and whether potential commonalities can guide monitoring, modelling, and conservation. Using a habitat modelling (CEPHALOPOD), we compare plankton diversity patterns across observation types and metrics, relating global open-ocean surface diversity from 330,000 observations of occurrence, abundance, biomass, and relative metagenomic reads to environmental climatologies. We find that diversity patterns converge on a latitudinal diversity gradient, with tropical oligotrophic gyres, western boundary currents and the Info-Pacific archipelago emerging as diversity hotspots. Occurrence-based estimates indicate higher diversity in eastern boundary upwellings than estimates accounting for community dominance, showing how multiple observation types provide complementary views of diversity. Surface ocean temperature, primary production, and physical transport emerge as remotely observable diversity proxies that can guide in situ sampling. Finally, diversity–ecosystem functioning relationships depend on the observation system: occurrence-based estimates highlight how diverse species can support production through complementary ecological roles, whereas abundance-, biomass-, and metagenomic-based estimates better reflect how dominant species contribute to processes such as carbon export. These findings can guide monitoring, modelling, and conservation under climate change.

## INTRODUCTION

Marine plankton diversity structures ocean ecosystems, and play a central role in global biogeochemical cycles (*1*, *2*). For example, phytoplankton growth drives 30 to 50% of global carbon fixation (*3*). Zooplankton transfer this fixed carbon to higher trophic levels via grazing, diurnal vertical migration, and the production of sinking fecal pellets, leading to particulate organic carbon flux to the deep ocean (*4*). As anthropogenic pressures increasingly alter marine ecosystems, plankton biodiversity monitoring needs to provide more scalable indicators of ecosystem functioning and resilience in support of conservation planning and ocean system modelling (*5*, *6*). In contrast to terrestrial ecosystems where biodiversity - ecosystem functioning (BEF) relationships have been extensively documented, such relationships remain poorly characterized in the global ocean (*7*, *8*). Large-scale terrestrial experiments have shown that higher taxonomic richness often indicates higher ecosystem productivity (*9*, *10*), stability and functional diversity (*11*). Whether these relationships translate to marine plankton, which span vastly different organisms and trophic levels, remains largely unknown. In particular, it remains unclear whether higher plankton diversity systematically promotes ecosystem functioning, which specific ecosystem properties are controlled by which facet of plankton diversity, and whether these relationships are linear in marine planktonic ecosystems (*8*). This knowledge gap arises because different *in situ* observation methods and diversity metrics capture only facets of diversity, such as a specific range of organism size, or treat community composition, structure, and relative dominance across taxa differently, thereby limiting our ability to resolve BEF relationships associated with marine plankton diversity (*12*, *13*).

In recent decades, traditional sampling methods such as net tows and microscopy have co-existed with newly emerging approaches, including quantitative imaging and molecular techniques, leading to a rapid increase in plankton *in situ* observations (*14*). Quantitative imaging has expanded data coverage to include fragile and previously under-sampled taxa, such as gelatinous zooplankton (*15*), while metabarcoding and metagenomics revealed cryptic diversity and functional potential across communities (*16*). Each *in situ* plankton sample captures only partial and method-or even instrument-specific snapshot of the local plankton community and diversity (*15*, *17*). On one hand, quantitative imaging and net-based observations are constrained by organism size and morphology and only cover eukaryotic phytoplankton and zooplankton (*15*, *18*). On the other hand, molecular approaches rely on marker genes with variable taxonomic resolution across groups, mostly used to cover prokaryotic plankton (*16*, *19*), but increasingly used to study eukaryotic plankton taxa (*20*, *21*). Beyond the observation type itself, diversity estimates also depend on how community structure is considered and weighted (*12*, *13*). While species richness only considers the number of taxa present, other metrics account for community evenness and penalize diversity estimates when the observed community is dominated by a few abundant taxa. Consequently, *in situ* observation types and diversity metrics each capture different components of community structure, and may therefore yield contrasting estimates of diversity (*13*). Determining where and when complementary facets of diversity converge in terms of latitudinal gradients, basin-scale features, or consensus hotspots is essential to develop robust diversity indicators that reflects ecosystem functioning and service provision to inform conservation planning and ocean modellers (*6*). For examples, regions with high diversity and substantial BEFs may be suitable in terms of the definition of MPAs as associated with the BBNJ agreement and the goal to conserve 30% of the ocean by 2030 (*22*). Diversity hotspots that are consistent across sampling and analysis methods, may also provide suitable testbeds for the development of transferable or method-specific indicators of plankton diversity that could support conservation planning even where only a subset of the plankton community has been observed. In contrast, identifying where complementary facets of diversity diverge is essential to identify where complementary types of observations are required to fully characterize plankton diversity, and if so, at which spatiotemporal resolution each method should be deployed to maximize knowledge gain while minimizing cost and effort.

Although they target different size classes, taxonomic groups, and ecosystem structure properties (*18*), broad-scale plankton diversity estimates may present converging patterns at climatological scales. This is because common environmental features, including temperature or light availability, that are latitudinally structured have been shown to consistently explain the distribution of marine taxa in the global ocean, from prokaryotes to megafauna (*23–25*). Many of these environmental features are routinely observed from satellites or global ocean reanalyses (*26*, *27*), raising the possibility of identifying environmental proxies from remote sensing, that consistently predict plankton diversity across complementary observation types and diversity metrics. However, *in situ* observations, or plankton diversity estimated from those, are spatially discrete, scarced and biased to well-observed regions such as the coastal ocean, which limits our ability to assess whether derived diversity patterns are representative for global ocean dynamics at multiple spatiotemporal scales (*19*, *28*).

Statistical habitat modelling addresses this limitation by empirically relating various types of biological observations to their associated environmental conditions, thereby enabling their spatial extrapolation into unsampled regions (*29*). Habitat models are particularly well suited to study plankton, as their physiology and ecological niches are tightly constrained by gradients of temperature, light, and nutrients or food availability (*30*, *31*). This approach, widely applied to marine ecosystems (*32*), provided global estimates of the spatial distribution of plankton biodiversity and its physico-chemical controls, from prokaryotes to zooplankton (*33–35*). Yet only 10% of habitat modelling studies include observations that account for community evenness (*32*) such as abundance, biomass or metagenomic reads. These are essential factors for studying the link between diversity and ecosystem properties (*36*), and, when taken into account, model outputs are rarely intercomparable (*37*). Even though recent studies have begun to link bacterial, phyto-, and zooplankton diversity to key ecosystem functions (*24*, *38*, *39*), most assessments still focus on species richness (*32*). Furthermore, diversity metrics are not direct proxies of ecosystem function. As a result, it remains unclear which aspects of diversity (e.g., richness, evenness) should be quantified to best capture the relationship between plankton diversity and each ecosystem property (*12*, *40*).

Ecological theory states that different ecosystem functions should depend on different facets of diversity (*13*). For example, Net Primary Production (NPP) is expected to benefit from the *niche complementarity* hypothesis, where higher species richness enhances ecosystem productivity through more efficient and complementary resource use across taxa, promoting the transfer of energy toward higher trophic levels by a broad fraction of plankton species (*24*, *41*). However, species richness strongly depends on sample size (i.e., volume and time range), as the large fraction of plankton taxa persisting at low abundance is detected differently across observation types and sampling efforts, which may lead to under-or overestimation of true plankton diversity (*42*, *43*). Some studies show that the *mass-ratio hypothesis* could help explain the role of plankton diversity and dominant species, or traits, in the processes of the biological carbon pump (*24*, *41*). For instance, particulate organic carbon export is driven by fast sinking fecal pellets from a subset of large zooplankton taxa that tend to dominate the biomass (*1*, *4*). Therefore, diversity metrics accounting for community evenness should enable to better reflect the role of species in regulating these ecosystem functions, as they allow to integrate dominance patterns (*44*). Yet, species richness remains the most widely used descriptor of biodiversity across marine studies, as its ease of measurement, and comparability across datasets make it a convenient proxy. As a result, richness is frequently used to study BEF, even though theory suggests that other facets of diversity may be more appropriate (*12*, *13*). Taken together, these challenges raise a broader question: can we identify global-scale plankton diversity indices, or environmental proxies, that transcend observation methods and diversity metrics to provide robust predictions of ecosystem functioning in space and time ?

To identify robust plankton diversity indices across *in situ* observation types and diversity metrics, we test four hypotheses. First, we test the hypothesis that the latitudinal diversity gradient (LDG) emerges as a consensus feature of marine plankton diversity across all observation strategies and diversity metrics. Second, we test the hypothesis that complementary *in situ* observation types and diversity metrics reveal systematic regional deviations from this consensus pattern because they emphasize different components of plankton community structure (e.g. rare versus dominant species). Third, we test the hypothesis that a reduced set of environmental drivers, such as temperature, is found as dominant drivers of multiple facets of plankton diversity, providing indicators for plankton diversity monitoring using remote sensing tools. Finally, we test the hypothesis that no single diversity metric universally explains BEF relationships, and that: (i) ecosystem functions governed niche complementarity are better explained by species richness; whereas (ii) functions dominated by a few abundant taxa are better explained by diversity metrics accounting for community evenness. To address these hypotheses, we leverage *in situ* plankton observations of occurrences, abundance, biomass and metagenomic reads from the AtlantECO (*18*) and the Microbiomics (*16*) data collections. We use CEPHALOPOD (*37*), a state-of-the-art habitat modeling ensemble framework designed to infer and intercompare plankton biodiversity estimates across observation methods and diversity metrics.

## RESULTS AND DISCUSSION

### 2.1. A consensus plankton diversity index to support conservation

Here, we first test whether the LDG is a consistent property of plankton communities retrieved across all combinations of *in situ* observation types and diversity metrics on the full global ocean. We then use this consensus diversity pattern to identify areas of interest for diversity conservation.

We retrieved global species-level plankton *in situ* observations from the AtlantECO (*18*) and Microbiomics (*16*) data collections. These include 21.1M occurrences, 19.7M abundances and biomasses, and 42.9M relative abundance of metagenomic reads (**Figure S1**). We estimated diversity using Hill numbers (*45*), that is the number of equally common taxa that would yield the observed diversity value, and allows to penalize rare taxa relative to dominant ones by increasing the scaling factor (*q*) (*42*).

Using CEPHALOPOD to generate robust global plankton diversity projections (**Figure S2** and **S3**; *see methods*), we show that the consensus diversity index averaged across combinations of *in situ* observation type (*n* = 4), Hill scaling factor (*n* = 21), month (*n* = 12) and uncertainty replicate (*n* = 10), presents a Spearman *p* of 0.39 with absolute latitude (p-value < 0.05; **Figure 1A**). The diversity index, that is rank normalized to values between 1 and 100, exceeds 75% in the tropical ocean (30°N to 30°S). In contrast, the diversity index drops below 30% in the high latitudes (beyond 50°N and S), with the Southern Ocean is identified as extrapolation outside environmental conditions of the observations (**Figure S3B**). This convergence towards a LDG is consistent with previous findings showing a decrease of species richness with latitude at the functional group level (*35*, *46*), and suggests that the LDG is also the predominating pattern when considering dominance patterns within the plankton community. We hypothesize that warmer low-latitude waters sustain higher plankton diversity because temperature influences physiological rates and allows more taxa with distinct thermal niches to coexist (*24*, *31*), while cooler high-latitude environments support fewer taxa due to harsher thermal constraints and shorter growing seasons (*47*). Likewise, this temperature effect also explains the seasonality of the LDG, with an increase in the diversity index of 20% between summer and winter months in subtropical (30°N and S) and high latitudes (above 50°N; **Figure 1B, S4**). Indeed above 45°N, warmer summer temperature associated with growing season support more diverse plankton assemblage than in winter (*35*, *48*)(**Figure S4**).

**Figure 1:**
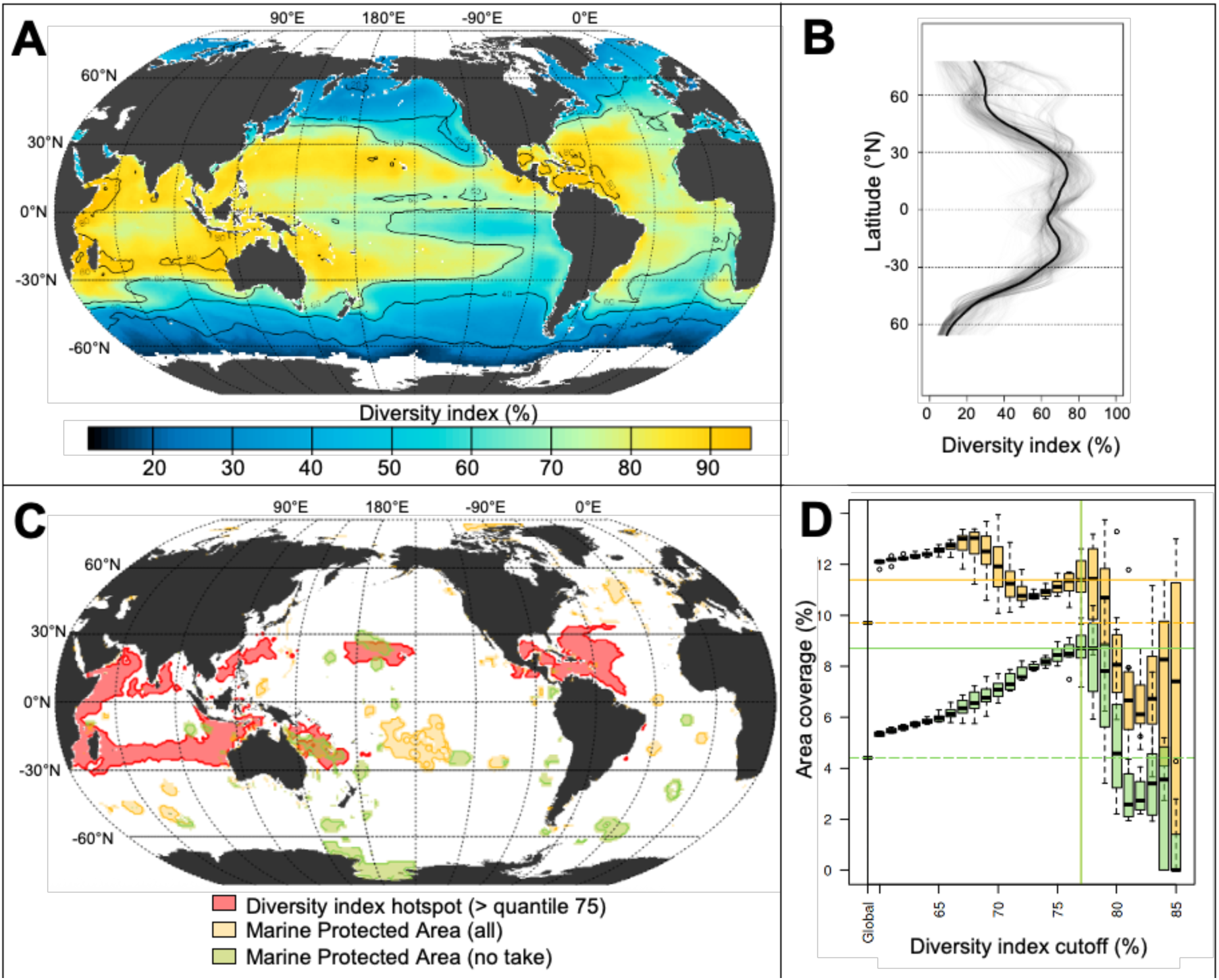
Consensus estimation of plankton diversity indices across all in situ observation types, month and hill scaling factor considered. Panel (**A**) present the consensus global diversity index pattern and (**B**) latitudinal profile of the consensus diversity index (in black) and all combinations of in situ observation types, month, and hill scaling factors (in grey). Fraction of the ocean identified as a diversity hotspot (above the 75th percentile) and the corresponding coverage by Marine Protected Areas (MPAs) under different protection categories (all; no take). **(D)** Percentage of each diversity index quantile covered by MPAs. Quantile 1 represents the global ocean and provides the baseline MPA coverage against which the coverage of higher diversity quantiles can be compared with.

The highest diversity index values (70 to 90%) occur in tropical western boundary systems, indicating that plankton diversity in these regions ranks within the top 10% to 30% of values across all combinations of *in situ* observations, Hill scaling factor, and months. Our consensus diversity index provides a robust assessment of whether the existing MPA network overlap with regions supporting high plankton diversity (**Figure 1C–D**). Globally, 11.5% of regions where the diversity index exceeds 75% overlap with MPAs, including 8.6% within no-take areas (**Figure 1D**). In contrast, MPAs cover 4.3 to 9.8% of global ocean, depending on protection level. However, this overlap is spatially uneven, with no overlap between diversity indices above 75% and MPAs in the Indian and Atlantic Oceans (**Figure 1C**). Some studies suggest that higher plankton diversity is associated with greater resource-use efficiency and increased resilience to shifts towards dominance by opportunistic bloom-forming taxa (*49*, *50*). Protecting regions and environmental conditions that sustain high plankton diversity may therefore help maintain plankton mediated ecosystem services including marine food-web productivity, carbon export, and biogeochemical cycling (*51*, *52*). Therefore, our consensus diversity index provides a complementary layer for conservation practitioners and policymakers to evaluate the value of MPAs for plankton biodiversity conservation, and incorporates this often overlooked component of ocean ecosystems (*51*, *52*).

### 2.2. Regional divergences in plankton diversity estimates

Then, we tested whether *in situ* observation types and diversity metrics explain systematic deviations from the LDG, and use these divergences to identify regions where complementary diversity indices are most needed to characterize the plankton community.

To isolate deviations from the LDG associated to observation types, we first computed, for each observation types, the mean diversity index averaged across hill scaling factors (*q*), months, and bootstrap replicates. We then quantified the effect of observation types on diversity indices by computing the standard deviation across observation types (*see methods*; **Figure 2A**). The largest deviations occur in the eastern tropical Pacific and Benguela upwelling system, as well as in the Kuroshio, California current and Gulf Stream system, where standard deviation in diversity index exceeds 30% (**Figure 2A**). This indicates that no single *in situ* observation type captures all facets of plankton diversity in these regions. To identify which observation type drives these regional differences, we compared latitudinal profiles (**Figure 2A**) and diversity index maps (**Figure S5**) for each observation types, averaged across all other factors. Only occurrence-based diversity indices present a unimodal latitudinal profile with values ranging from 60 to 70% between 35 and-35°N (**Figure 2B**), and regional maxima coinciding with the highest observational disagreement (**Figure 2A**, **S5A**). Due to their relative ease of measurement, occurrences capture broad taxonomic richness without accounting for community structure (*12*, *24*), highlighting the large number of co-occurring taxa in these regions (*53*, *54*). In contrast, abundance, biomass and metagenomic observations resolve community dominance and evenness, and associated diversity indices are penalized when a few taxa dominate the plankton community, such as in highly productive environments (*55*). Therefore, abundance-, biomass-, and metagenomic-based diversity indices present similar bimodal latitudinal profiles and spatial distributions centered on tropical oligotrophic gyres (**Figure 2B, S5**). The latter coincides with regions where the standard deviation in diversity index ranges between 0 and 10 %, revealing consistent estimates in these areas (**Figure 2A**). Together, these results identify eastern boundary upwelling systems and western boundary currents as priority areas to enforce complementary observation types, because no single approach resolves both broad taxonomic richness and community structure (*12*, *13*, *36*). Therefore, our maps provide a practical basis for observing networks to prioritize complementary sampling efforts, while improving plankton diversity assessments used by conservation practitioners and ocean modellers.

**Figure 2:**
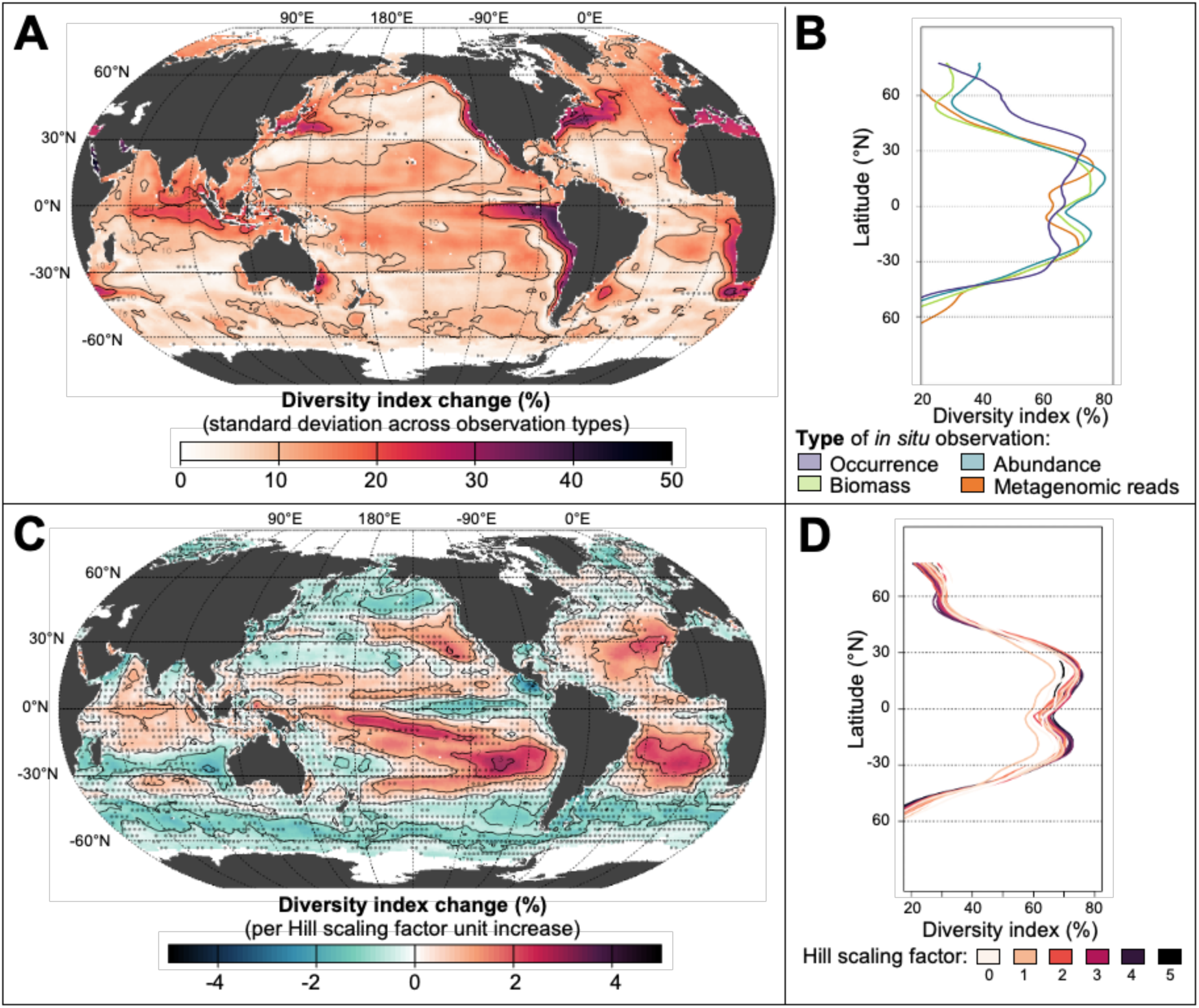
Variance in plankton diversity with (**A**) the diversity index change (%) expressed as the standard deviation between each observation types across all unique combination of hill scaling factors, months, and bootstrap replicates. Panel (**B**) shows the latitudinal profile associated to each observation types, averaged across all other factors. Variance in plankton diversity with (**C**) the diversity index change (%) expressed as stepwise increase for one Hill scaling factor unit (e.g., from *q* = 0 to *q* = 1) across all unique combination of observation types, months, and bootstrap replicates. The (**D**) panels show the latitudinal profile associated to each hill scaling factors, averaged across all other factors.

To isolate the deviations from the LDG associated to diversity metrics, we computed, for each Hill scaling factor (*q*), the mean diversity index averaged across observation types, months, and bootstrap replicates. We then quantified the effect of increased penalization of the diversity index when community composition is uneven, using a moving window of Δ*q* = 1 (**Figure 2C**). The largest negative deviations occur in eastern tropical upwelling systems, and temperate to high latitudes, above 40°N and 40°S, where the diversity index decreases by 2% for each unit increase in *q* (**Figure 2C**). This indicates that accounting for community evenness consistently lowers diversity indices in these regions, suggesting that plankton communities are dominated by a few abundant taxa (*12*, *42*). Indeed, highly productive environments, such as the eastern Pacific subtropical upwelling, are characterized by high phytoplankton abundance (*53*, *54*), whereas low-productivity high-latitude regions are dominated by macrozooplankton taxa like krill (*56*). Conversely, the largest positive deviations occur along the eastern margins of the subtropical gyres, where the diversity index increases by 2% for each unit increase in *q* (**Figure 2C**) and up to 4% for abundance-and metagenomic-based estimates (**Figure S6B** and **D**). Rather than being dominated by a few abundant taxa, subtropical gyres support relatively even plankton comunities all year round, maintained by stable environmental conditions (*57*, *58*) (**Figure 2C**). Consequently, diversity estimates in these regions are not penalized when greater weight is given to community evenness. Instead, their relative importance increases as diversity indices decrease in productive and high-latitude regions, shifting diversity maxima towards the subtropical gyres. The latitudinal profiles of diversity indices support this result as the amplitude of the bimodal pattern increases with higher Hill scaling factor (**Figure 2D**). Although the influence of diversity metrics is weaker than that of observation type (**Figure 2A** and **C**), varying the Hill scaling factor provides modellers with an efficient way to resolve large-scale gradients in community evenness without requiring complementary observing systems (*12*, *42*).

The systematic regional deviations between highly productive regions, dominated by a few abundant taxa, and low-productivity high-evenness regions are consistent facets of diversity recognized in both terrestrial and marine ecosystems (*13*, *23*, *59*). This also suggests that environmental drivers beyond the broad latitudinal temperature gradient contribute to structuring different facets of plankton diversity (*21*, *25*), motivating the search for environmental predictors that consistently explain these complementary patterns.

### 2.3. Environmental drivers of diversity

Next, we tested whether plankton diversity patterns, the divergences across *in situ* observation types and diversity metrics, can be explained by a reduced set of environmental drivers, thereby enabling the development of remote sensing proxies for large-scale diversity monitoring.

To quantify and rank the environmental features that govern the spatial distribution of our diversity indices, we performed a redundancy analysis (RDA; *60*) of these features against all diversity index estimates at a global scale (**Figure 3A**). The RDA identifies the major axes of variation in diversity indices that are constrained by linear combinations of environmental features, thereby revealing which environmental feature best explain diversity indices. We show that each diversity index is positively associated with RDA1 (62%; **Figure 3A**). The first order gradient of all diversity indices (RDA1) is strongly associated with temperature (0.95), opposite to nutrients (-0.88), dissolved oxygen (-0.92), and Dissolved Inorganic Carbon (DIC; - 0.76) concentrations (**Figure S7**), all of which are latitudinally structured. These environmental variables correspond to the features explaining most variance in the algorithms composing the model ensembles (50 to 70 %; **Figure S8**, cluster 2 and 3). This shows that temperature is a proxy of plankton diversity indices, following the LDG and observable from space (**Figure 3**; **S7**)(*26*). This is consistent with the literature, as warmer low-latitude waters sustain higher species richness because temperature influences physiological rates and allows more taxa with distinct thermal niches to coexist (*24*, *31*), while cooler high-latitude environments support fewer species due to harsher thermal constraints and shorter growing seasons (*47*)(**Figure 3B**, *left panels*).

**Figure 3:**
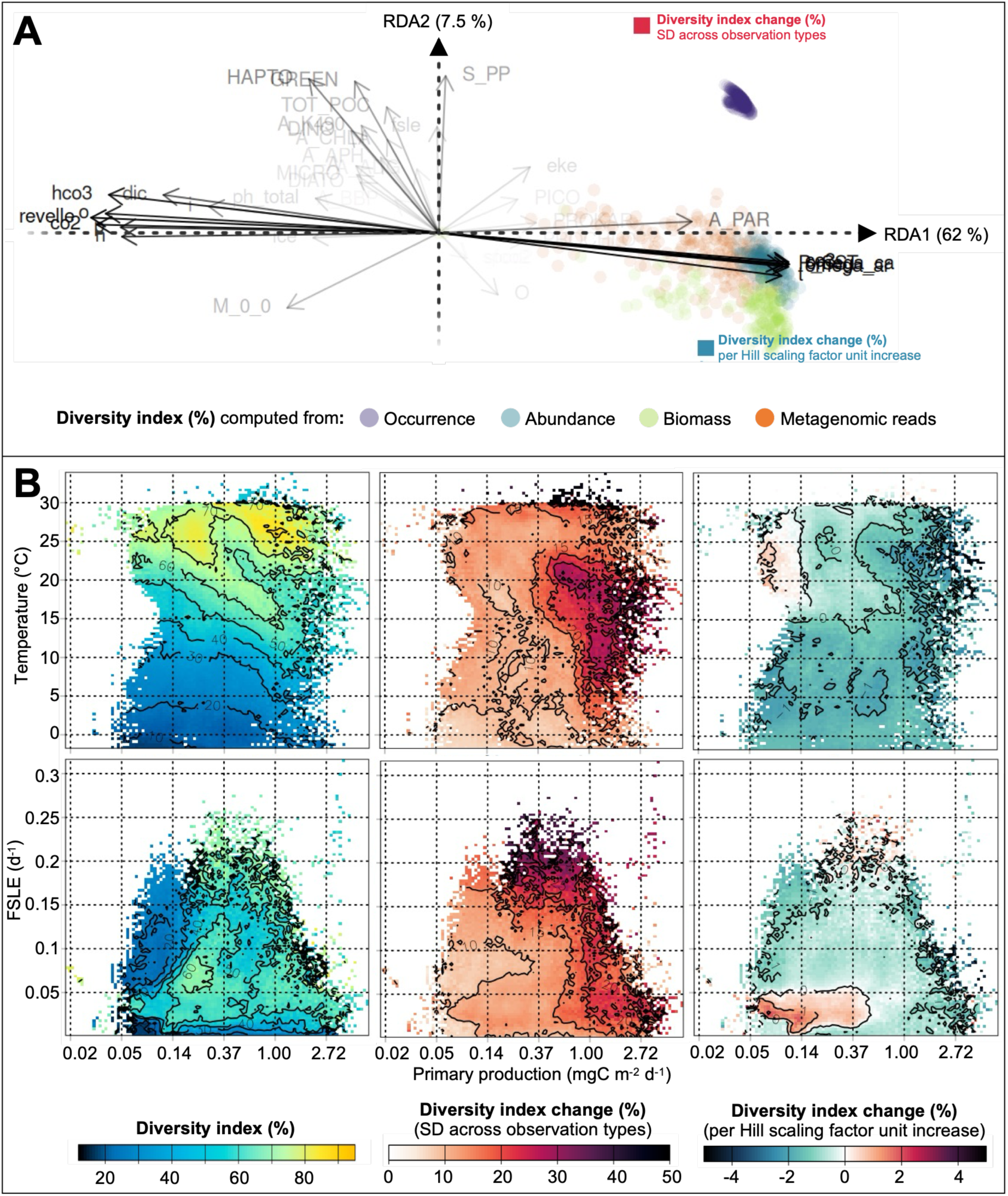
Environmental drivers of plankton diversity. Panel (**A**) shows the Redundancy Analysis (RDA) relating projected diversity indices across *in situ* observation types and Hill scaling factors to environmental features. Diversity indices are colored by observation types. The standard deviation in diversity index across *in situ* observation types and change in diversity index for one unit increase in the Hill scaling factor are represented in red and blue, respectively. For readability, only abbreviated labels are displayed for environmental features in the RDA space; corresponding full names are provided in **Table S1**. Panel (**B**) shows biplots in the environmental space defined by temperature, primary production, and Finite Size Lyapunov Exponent (FSLE), corresponding to proxies of RDA axes 1, 2, and 3, respectively. The biplots display the average diversity index, the standard deviation in diversity index across observation types, and the diversity index change associated with increasing the Hill scaling factor by one unit in the left, middle, and right panels, respectively.

The diversity index change across *in situ* observation types associates to RDA2 (0.75; **Figure 3A**). This second RDA axis is associated with primary production (0.57), mixed layer depth (-0.27) and oxygen saturation (-0.23; **Figure S7**). These features were also found as explaining only 30 to 50% of the variance in the algorithms composing the model ensembles (i.e., cluster 1 and 4; **Figure S8**). Diversity index change above 30% occur in highly productive (> 1 mgC m^-2^ d^-1^) or high mixing conditions (FLSE > 0.15 d^-1^; **Figure 3B**, *middle panels*). Indeed, we show that occurrence-based diversity indices are highest in productive, low-latitude regions such as coastal upwelling systems (**Figure S5A** and **S9C**), where strong mesoscale activity, nutrient enrichment, and environmental heterogeneity (*31*, *55*) promote the co-existence of species (*31*, *61*). Therefore, occurrence-based diversity indices reveal regions where many species are present due to niche overlap (*54*, *62*), but this does not imply that all species are evenly contributing to community composition or ecosystem processes as this observation type does not resolve dominance patterns. In contrast, abundance-, biomass-and metagenomic-based indices show that productive regions display lower diversity, as a few dominants often outcompete other taxa for resources and drive ecosystem functioning. Therefore, in addition to temperature, primary production (i.e., derived from Chlorophyll-a; GMIS) is a remote sensing proxy discriminating co-existence from evenness driven plankton communities.

The diversity index change per Hill scaling factor unit increase also associates to RDA2 (-0.42; **Figure 2C, 3A**). Positive values, meaning that the diversity index increases when greater weight is given to community evenness through higher Hill scaling factor, are found in low productivity (< 0.4 mgC m^-2^ d^-1^) and low mixing (FLSE < 0.05 d^-1^; **Figure 3B**, *right panels*). This relationship between diversity index, community evenness, and productivity is associated to temperatures above 15°C (**Figure 3B**, top right panel). In the tropical ocean, increasing the Hill scaling factor penalizes high mixing - high productive regions such as the eastern Pacific tropical upwelling (**Figure 2C**), where a few species are known to dominate the plankton community (*53*, *54*). In contrast, increasing the Hill scaling factor increases the diversity index in conditions of low mixing and intermediate productivity (*63*, *64*) that typically occur at the boundaries between oligotrophic gyres and upwelling systems, where both apparent species richness and evenness remains high (*33*, *57*)(**Figure 2C** and **3B**). Therefore, primary production (derived from chlorophyll-*a*; GMIS)(*27*) and FSLE (derived from altimetry-based geostrophic velocities)(*65*) emerge as two satellite-observable proxies of plankton community evenness in the tropical ocean.

Together, these results are consistent with previous literature showing that temperature dominates large-scale climatological gradients in plankton diversity (*24*, *33*), whereas mechanistic models suggest that mesoscale activity (e.g., FSLE) contributes to regional diversity patterns by increasing environmental heterogeneity, species dispersal, and opportunities for species coexistence (*66*), thereby influencing apparent richness and community structure (*67*). Together with primary production, these variables provide the remote sensing community with diversity proxies to enforce ocean monitoring across facets of plankton diversity and scales, and may help reconcile the different perspectives of statistical and mechanistic approaches in predicting plankton diversity (*24*, *66*).

### 2.4. Diversity to ecosystem properties relationships

Finally, we assessed whether relationships between global plankton diversity and ecosystem properties are consistent across *in situ* observation types (Figure 4), and whether the divergences across observation types translate into predictable differences in BEF relationships. This is essential for ocean and ecosystem modellers to help them determine which diversity index best predicts which ecosystem function. Our findings show that, while the overall shape of BEF relationships is largely insensitive to the choice of *in situ* observation type, their magnitude varies substantially among observation types at the global scale.

**Figure 4:**
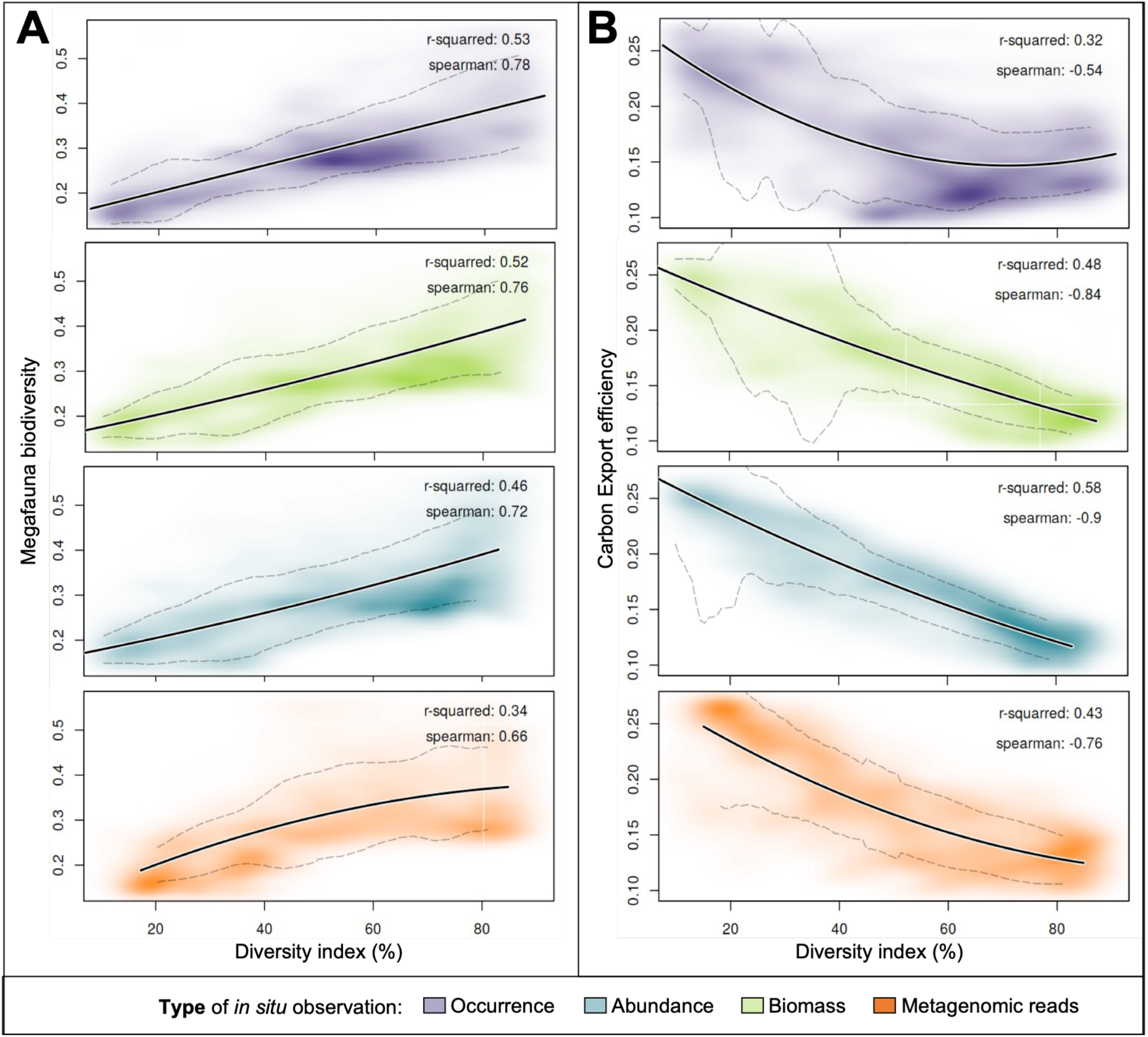
Emergent relationships between diversity index (%) and ecosystem properties or functions for each *in situ* observation types, with (A) Global oceanic megafauna biodiversity expressed as normalized species richness and (B) the efficiency of the production exported (POC/NPP; no units). The strength of the shading represents the density of geographical cells associated to a diversity index and ecosystem property bin in the global ocean. The continuous and dashed lines correspond to the average ± 1 standard deviation of a second order polynomial regression of the diversity index against the ecosystem property. The R^2^ represent the goodness of fit of the regression while the spearman correlation indicates the strength of the linear trend.

We provide evidence that plankton diversity correlates with other compartments of the marine ecosystem such as megafauna diversity (*68*)(**Figure S9A**). This is expected from previous studies based on species richness (*24*), and common environmental drivers shaping marine diversity across trophic levels (*25*). We found a positive Spearman correlation (*p* = 0.73 ± 0.05; linear model accuracy R^2^ = 0.46 ± 0.09) between global megafauna diversity (*68*) and plankton diversity indices across all observation types (**Figure 4A**). Above diversity indices of 80 %, megafauna diversity values range between 0.3 and 0.5, indicating that both variables present overlapping hotspots. The relationship between plankton diversity and megafauna diversity is weaker for metagenomic-based diversity estimates (*p* = 0.66; R^2^ = 0.34). This was expected since this observation type targets prokaryotes (*16*), that are the most indirectly connected to marine megafauna in the context of trophic interactions. These findings suggest coherent trophic structuring from the base of the food web to apex predators (*31*, *68*). This pattern is ecologically consistent, as fish and marine mammal distributions depend on similar environmental drivers (*69*) as well as on the abundance, composition, and seasonality of their planktonic preys (*31*, *70*).

Second, we provide evidence that specific diversity indices correlate with specific components of the biological carbon pump (**Figure S9B** to **D**). This is expected as the effect of diversity on the efficiency of carbon export within the biological carbon pump follows a *mass-ratio* hypothesis (i.e., driven by dominant taxa) while the role of diversity on primary production follows a *niche complementarity* hypothesis (i.e., driven by the complementary ecological role of all taxa present)(*24*, *41*). We show a negative Spearman correlation (*p* = 0.76 ± 0.16; R^2^ = 0.45 ± 0.10) between the efficiency of carbon export within the biological carbon pump, expressed as the share net primary production (NPP) exported to the deep ocean layers (*64*) and plankton diversity indices across all *in situ* observation types (**Figure 4B**). Export efficiency is primarily driven by large diatoms and large macrozooplankton swarms, especially at high latitudes (*64*, *71*). In lower latitudes and oligotrophic gyres, more functionally balanced communities with diverse sizes and trophic strategies enhance carbon use and bacterial remineralization, thus lowering the fraction of NPP exported. Consistent with this mechanism, occurrence-based diversity indices, that do not resolve community structure, exhibit a weaker relationship with carbon export efficiency (*p* =-0.54; R^2^ = 0.32). In contrast, abundance-, biomass-or metagenomic-based diversity indices that account for plankton community evenness are better to analyze the efficiency of carbon transfer within marine ecosystems (*39*, *64*) and to be considered in ocean and ecosystem models (*6*). Conversely, occurrence-based diversity indices better capture emergent relationships with other aspects of the biological carbon pump (*64*), such as carbon export flux (*p* = 0.37; R^2^ = 0.08; **Figure S10A**) and, as expected from **Figure 3B**, net primary production (*p* = 0.67; R^2^ = 0.35; **Figure S10B**). These emergent relationships are consistent with the *niche complementarity* hypothesis (*24*, *41*), whereby a wide range of taxa contribute to net primary production and carbon export fluxes (*1*, *8*).

Together, these results support our hypothesis that no single facet of plankton diversity can universally capture all BEF relationships. Instead, the most informative diversity index depends on the ecosystem property considered: richness-based indices better describe functions emerging from species complementarity in ressource use, whereas abundance-, biomass-, and metagenomic-based indices that account for community dominance better capture functions governed by the mass-ratio hypothesis. Selecting diversity indicators according to the ecological process of interest should therefore improve the representation of marine ecosystem functioning in monitoring programs and ocean models (*6*).

### 2.5. Caveats and limitations

In this work, we showed how the LDG as a consistent pattern of plankton diversity across observation types and diversity metrics. We also showed that temperature, primary production and physical transport are three satellite-observable proxies that explain the LDG, as well as regional deviations from the LDG (**Figures 1** to **3**). Because these conclusions rely on integrating complementary *in situ* observations (*16*, *18*), it is important to consider how sampling and taxonomic limitations may affect their representation of plankton diversity (*28*, *72*).

First, incomplete coverage of the plankton community limits our ability to disentangle taxonomic and observation-type effects. Each *in situ* observation type captures only a subset of the plankton community. Traditional net sampling used for occurrences, abundances and biomass observations, is known to under-sample small organisms or fragile taxa because of mesh-size selectivity (*28*, *73*), whereas metagenomic approaches primarily characterize prokaryotic communities (*16*, *17*, *74*)(**Figure S1C**). Consequently, differences among diversity indices may partly reflect the taxa accessible to each observation type, in addition to differences in community structure. Second, taxonomic resolution also differs among observation methods. Many organisms cannot be identified reliably to species level because of body damage during collection or limited morphological resolution, particularly for abundance and biomass observations (*15*, *75*). Quantitative imaging avoids destructive sampling but provides lower taxonomic resolution (*76*). This may lead to the overrepresentation of certain plankton groups as copepods (*77*), thereby affecting the attribution of local richness estimates to the diversity of certain taxa differently across observation types. However, these limitations do not affect the consistent identification of the LDG, and importantly, reflect the biases encountered by observers, or modelers and policy makers (*28*). Our conclusions therefore reflect the level of information available to these stakeholders when assessing, predicting, and managing ocean biodiversity.

Finally, our conclusions apply to the plankton community level but do not indicate whether the LDG, and its associated environmental drivers are conserved within individual taxonomic groups. This was previously shown for occurrence-based estimates across phyto-and zooplankton (*35*), but not for prokaryotes (*46*). For abundance and biomass observations, species-level taxonomic resolution was insufficient to generate reliable species-specific projections (*18*, *75*). For metagenomic observations, relative abundances cannot be summed across species-level projections because each sample is normalized to a total of one (*16*). We therefore projected community-level diversity directly (*78*), which allows robust comparison across observation types but prevents us from identifying which taxa dominate each plankton community or testing whether the LDG is consistent across individual taxonomic groups.

### 2.6. Perspectives

Our results provide guidance on where and when complementary observations are most needed. Rather than increasing the number of observations globally in an opportunistic way, future observing efforts combining complementary observation types should systematically target areas where diversity estimates diverge substantially between observation types, or fluctuates through the year, such as in eastern boundary upwelling systems and western boundary currents **(Figure 2**). Long-term monitoring and time series (e.g., ALOHA) serve a complementary objective, providing essential insights into temporal and climate-driven changes in biodiversity rather than being designed primarily to characterize global patterns of diversity. Moreover, as abundance-, biomass-and metagenomic-based estimates provide similar global diversity patterns, current developments in metagenomic approaches targeting universal marker genes (*79*) and omics-derived abundance estimates (*80*) hold potential for a more standardized and potentially less costly observing strategies.

Our identification of satellite-observable environmental drivers of plankton diversity (**Figure 3**) confirms that diversity patterns are linked to environmental variables routinely monitored from space, suggesting that machine-learning algorithms could use these variables to predict global plankton diversity and guide where additional in situ observations would be most informative (*81*, *82*). Temperature, productivity and variables associated with physical transport such as Finite Size Lyapunov Exponent (FSLE) could be used to identify regions where specific diversity facets are either changing rapidly or most likely to diverge, and thus where additional *in situ* observations provide the greatest added value to remote sensing. Testing the transferability of these relationships across spatial and temporal scales will be essential, particularly in highly dynamic environments where mesoscale processes alter plankton community composition and consequently BEFs relationships (*66*, *83*). Satellite-based estimates of diversity changes may also be useful to monitor the response of marine plankton ecosystems to climate extremes such as marine heatwaves or to define dynamic marine protected areas (*22*).

Our results also provide a basis for extending these recommendations beyond taxonomic diversity to functional and trait-based diversity, both of which have been associated with aspects of ecosystem functionning such as carbon export (*1*, *4*) or productivity (*8*, *13*). These dimensions are currently largely based on occurrence observations (*41*, *84*), but the emergence of metagenomic approaches targeting functional genes, provides an opportunity to test whether the observation-type effects identified for taxonomic diversity also apply to functional diversity (*20*, *34*).

Furthermore, the new imaging methods provide unprecedented insights into global plankton trait biogeography, and are increasingly deployed on autonomous platforms, including gliders and floats within the Bio-Argo fleet, they offer promising opportunities to expand spatial coverage and complement current global plankton observations (*15*). Determining how observation type and diversity metric influence estimates of functional diversity and their relationships with ecosystem processes will alo be important for improving marine ecosystem and biogeochemical models, for example those used in the IPCC to project future changes in ecosystem structure and function (*6*).

## 3. Conclusion

Despite differences in how marine plankton communities are observed and diversity quantified, our study identifies robust commonalities across these estimates, that provide a baseline for large-scale diversity monitoring and conservation. The LDG is robust across these approaches, revealing a shared global spatial pattern that can inform comparisons across observation systems and help identify priority regions for conservation, including MPA planning. In contrast, eastern boundary upwelling systems and western boundary currents show the strongest divergence among diversity estimates, identifying regions where complementary observation strategies reveal different facets of plankton communities. These differences highlight the importance of matchin observation strategies to regional ecological conditions, rather than relying on a single approach globally. These divergences are associated with environmental regimes defined by temperature, productivity and physical transport, all of which are observable from satellites, and therefore can be used to diagnose where and when additional in situ observations may provide the greatest added value. Finally, the different facets of diversity shape emergent BEF relationships in a different way. Species richness is more closely associated with functions such as primary production and high trophic-level diversity, consistent with the idea that different species contribute complementary ecological roles (niche complementarity)(*24*, *41*). By contrast, metrics accounting for the relative dominance of common versus rare species govern ecosystem properties related to carbon export to the deep ocean, consistent with the mass-ratio hypothesis, whereby the most abundant organisms exert a disproportionate influence on ecosystem functioning (*24*, *41*). Together, these results show that plankton biodiversity cannot be adequately represented by a single observation strategy, which means that neither omics-based nor quantitative imagine based methods alone will capture the full spectrum of plankton diversity and its links to ecosystem functioning. Rather, the commonalities and divergences among approaches can be exploited to improve where and how diversity is monitored and linked to ecosystem functioning, which will also improve future projections of potential changes associated with climate change.

## METHODS

### 3.1. Data

#### 3.1.1. Occurrence

We retrieved plankton occurrences synthetized within the AtlantECO-BASE v2 dataset (*18*)(**Figure 5**, step 1). Out of the 21.1M observations, the dataset comprises 739,000 unique samples across 1,693 accepted phytoplankton and 1,540 accepted zooplankton species. Taxonomic annotation follows the World Register of Marine Species (WoRMS). The 477,000 unique samples collected by the Continuous Plankton Recorder (CPR) were excluded to avoid an overwhelmingly large spatial clustering of observations as they are geographically constrained to the north Atlantic and Southern Ocean, and between 7 and 10m depth (**Figure 5**, step 2). To reduce sampling heterogeneity, we regridded occurrences to a 1 x 1° spatial resolution, since 1900, and integrated over 0 to 50 m. We considered only species with at least 50 occurrences, corresponding to the minimum requirements for habitat modelling with CEPHALOPOD (*37*). To compensate the lack of true absences, we sample pseudo-absences following (i) the nearby density of presences (i.e., geographical bias equal to presences)(*85*) and (ii) in geographical areas characterized by environmental conditions outside those of the occurrences (i.e., niche-based sampling)(*37*). To minimize prediction variance, we sampled pseudo-absences in equal number and weight to the presences (D-design theory)(*86*) (**Figure 5**, step 5).

**Figure 5:**
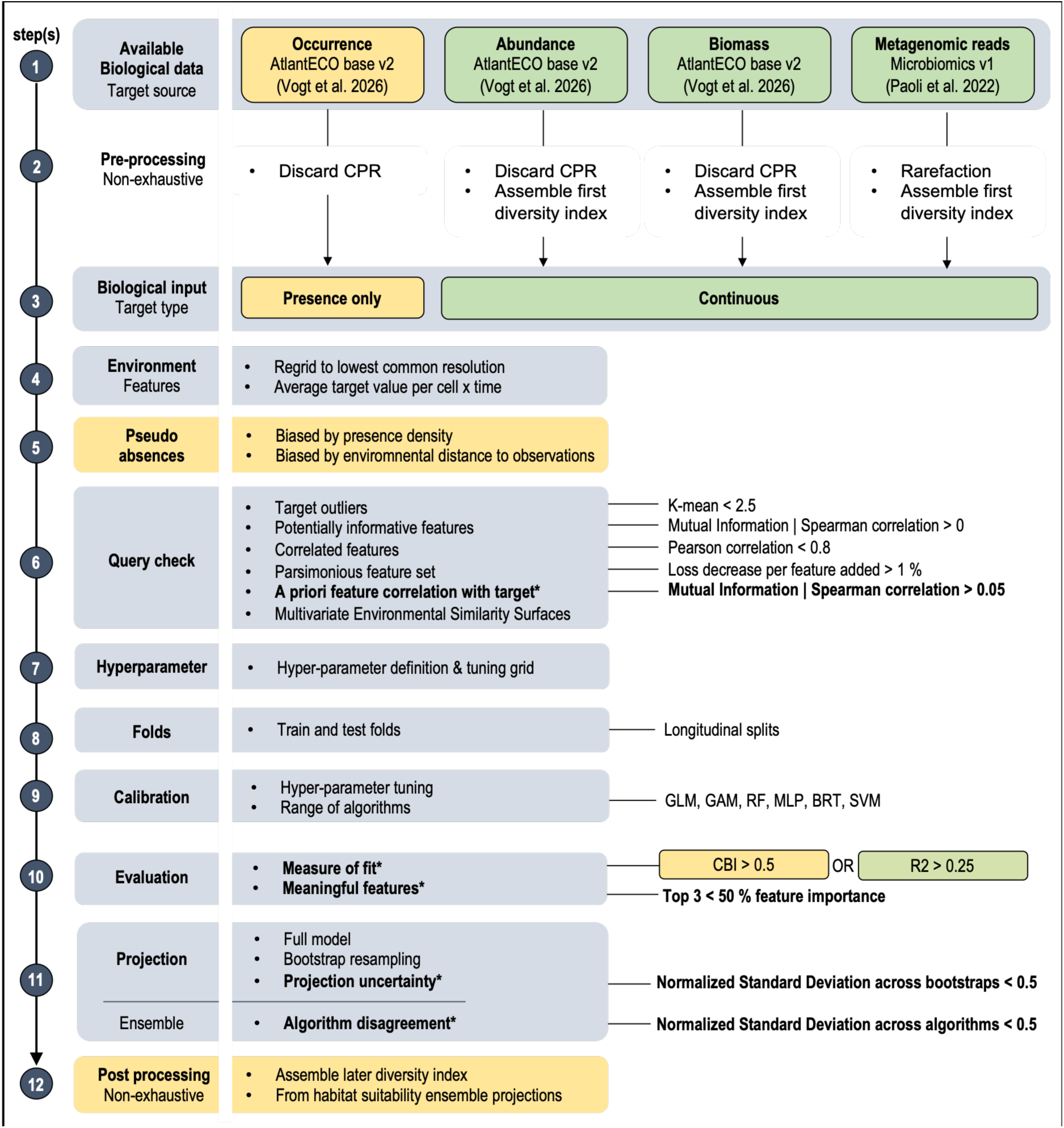
Synthetic diagram of the CEPHALOPOD framework, including pre-and post-processing steps of biological datasets required for the predict first (occurrence data) and assemble first (abundance, biomass and metagenomic data) diversity estimates. The main steps are indicated on the left-hand side. The associated quality checks or modelling choices are displayed in the middle panels. The corresponding thresholds used for quality flags are indicated on the right-hand side. Those used in the final traffic-light quality flag are indicated in bold. Adapted from Schickele et al. (2025).

#### 3.1.2. Abundance and biomass

We also retrieved the species-level observations of plankton abundances and biomasses synthetized in the AtlantECO-BASE v2 dataset (*18*) (**Figure 5**, step 1). Measured abundances were converted in biomass using individual carbon content estimates for each taxon (*18*). Our of the 19.7M observations, this dataset contains 652,000 unique samples across 1,884 accepted phytoplankton and zooplankton species. As for the occurrence dataset, taxonomic annotations follow those of the WoRMS. Again, the 548,000 samples stemming from the CPR were discarded. (**Figure 5**, step 2). To reduce sampling heterogeneity, we averaged abundance and biomass samples on the same spatial and temporal grid as for the occurrence dataset, and considered only species with at least 50 abundance or biomass observations in subsequent steps (*37*).

#### 3.1.3. Metagenomics

We retrieved plankton metagenomic reads from the Microbiomics v1 database (*16*) (**Figure 5**, step 1) at the mOTUs level, representing species-level clusters identified from multiple marker genes and annotated using the Genome Taxonomy Database (GTDB)(*74*). Out of 45.9M observations, the dataset comprises 308 unique samples, encompassing 18,707 mOTUs, associated with 4,016 known taxa. Observed richness ranged from 183 to 2676 mOTUs (Q25: 825; median: 1,105; Q75: 1,388). To limit biases from uneven sequencing depth or variable genomic content, we rarefied samples to 1000 mOTUs, balancing reduced sample bias risk and representative community diversity (**Figure 5**, step 2)(*46*). To reduce sampling heterogeneity, we averaged metagenomic reads on the same spatial and temporal grid as for the other datasets, and considered only species present in at least 50 samples (*37*).

#### 3.1.4. Environmental feature collection

We used global habitat modelling to derive continuous climatologies (i.e., diversity indices or habitat suitability indices; see *section 3.2*. below) from spatially discrete *in situ* observations (*29*, *37*). We selected 47 monthly, observation-based environmental features at a 1° x 1° global resolution (**Figure 5**, step 4; **Table S1**) (*37*). These features describe the physical (e.g., sea surface temperature), chemical (e.g., nutrient concentrations, carbonate chemistry), and biological (e.g., chlorophyll-a concentration, primary production) properties of epipelagic waters, as well as circulation and turbulence indicators such as eddy kinetic energy (EKE). Vertically resolved environmental fields were integrated over the top 50m to match biological observations filtering.

#### 3.1.5. Ecosystem properties

To investigate BEF relationships, we compiled four ecosystem properties that are proxies of key plankton roles in carbon cycling and energy transfer to the upper trophic levels. Marine biodiversity of high trophic levels was represented by normalized global species richness of bony fishes, sharks, cetaceans, and squids (*68*). Plankton contribution to the ocean carbon cycle was represented by three facets of the biological carbon pump: carbon fixation with the surface NPP (mgC.m^-2^.d^-1^), pump intensity with particulate organic carbon export flux (FPOC; mgC.m^-2^.d^-1^) and pump efficiency with the e-ratio (FPOC/NPP;-) (*64*). We regridded them to the same 1 x 1° global grid to facilitate the comparison of these ecosystem properties to our plankton diversity indices.

### 3.2. Pre-processing for habitat modelling

To estimate diversity from species-level observations, two main approaches exist. The *predict first – assemble later* strategy starts by building species-level distributions and then derives diversity indices from these distributions. Conversely, the *assemble first – predict later* strategy computes diversity indices at sampling locations prior to modelling their distribution. The latter generally minimizes projection biases (*78*). As occurrence data are qualitative, only richness can be computed using the *assemble first - predict later* method. However, occurrences offer sufficient spatial coverage to estimate species-level distributions using CEPHALOPOD. Therefore, we generate species-level habitat suitability maps and derive multiple diversity indices following the predict first - assemble later approach (**Figure 5**, step 2). For abundance and biomass observations, we constructed pseudo-samples at the climatological scale by aggregating point observations at a 1° x 1°, monthly, spatio-temporal resolution, matching the resolution of environmental features. In contrast, abundance, biomass, and metagenomic reads datasets are quantitative but spatially sparse. Therefore, we computed diversity indices from in situ observations and then modeled their distributions using the assemble first - predict later approach (**Figure 5**, step 2). To ensure comparability across datasets, and account for differences in sensitivity to rare or dominant taxa, we use the hill diversity framework (*42*). This framework overcomes these limitations by expressing all diversity indices as Hill numbers, where diversity is expressed as the effective number of species (*12*, *42*). We considered Hill scaling factors (*q*) from 0 to 5, in 0.25 increments, encompassing species richness (*q* = 0), Shannon (*q* = 1) and Simpson (*q* = 2) indices.

### 3.3. Habitat modelling framework

To model species distributions from occurrence data and Hill diversity indices from abundance, biomass, and metagenomic data, we used CEPHALOPOD, a unified framework for habitat modeling across diverse marine datasets (*37*). First, CEPHALOPOD applies standardized pre-processing and quality control to biological targets and environmental features (**Figure 5**, step 6). To reduce model complexity, CEPHALOPOD constructs a subset of non-collinear, informative, and non-redundant features by discarding uninformative environmental features, computing pairwise correlation at the target locations, and performing Recursive Feature Elimination (RFE) (*37*, *87*). To identify regions where algorithm extrapolate into novel environmental space, CEPHALOPOD computes a Multivariate Environmental Similarity Surfaces (MESS) metric (*88*). This results in a curated set of input data and parameters suitable for habitat model calibration. Next, CEPHALOPOD implements an ensemble modeling approach combining multiple algorithms to reduce uncertainty linked to the model choice (*37*). To mitigate spatial autocorrelation, that may bias model estimates, and optimize algorithm and hyperparameter training, CEPHALOPOD performs a Spatial Block Cross-Validation (SBCV; **Figure 5**, steps 7 to 9). This balances spatial independence and data partitioning for algorithm training and evaluation (*89*). Algorithm performance is evaluated by comparing predicted and observed values for each evaluation fold (**Figure 5**, step 10). Furthermore, we compute the importance of each feature in explaining these observations (*37*). Both metrics are linked to quality flags (*see Section* 2.4.). Finally, CEPHALOPOD propagates uncertainty to the projections by running bootstrap replicates across all retained algorithms and taxa (**Figure 5**, step 11). A final quality check ensures that only robust and interpretable outputs are retained (*37*).

### 3.4. Quality assessment within CEPHALOPOD

To guarantee both robustness and interpretability, CEPHALOPOD applies a four-step quality control procedure. First, we test the relevance of environmental features by comparing their average Spearman correlation and mutual information with the biological target, against a null distribution of permuted target values (**Figure 5**, step 6). Environmental features are retained only if their combined signal exceeds the null expectation by at least 0.05 (*37*). Second, we evaluate predictive performance using SBCV folds (**Figure 5**, step 9). Algorithm skill is quantified with the Continuous Boyce Index for presence-only data or R² for continuous data, with minimum thresholds of 0.5 and 0.25, respectively (*37*). Third, we assess ecological interpretability based on the feature importance (**Figure 5**, step 10). Algorithms are considered interpretable when the three most influential features explain at least 50% of the target variance (*37*). Fourth, we quantify projection uncertainty using the Normalized Standard Deviation (NSD; **Figure 5**, step 11), defined as the standard deviation across bootstrap replicates normalized by the mean across all grid cells. Algorithms are retained only if their global mean NSD remains below 0.5 (*37*). Only algorithms meeting all four criteria are included in the species-level ensemble projections for occurrences, and diversity indices projections for abundance, biomass, and metagenomics.

### 3.5. From CEPHALOPOD outputs to mapped diversity estimates

To generate global diversity maps based on occurrence data and the *predict first – assemble later* strategy for occurrence data, we estimated diversity across the projected species habitat suitability at each geographical cell, using the same Hill scaling factors as for the other datasets (*q* = 0 to 5, by 0.25; **Figure 5**, step 12). At this stage, we produced diversity maps for 4 datasets, 12 months, 10 bootstraps and 21 Hill scaling factors, capturing variations in observation types, community evenness weighting, resulting in 10,080 spatial projections. To enable quantitative intercomparison of diversity across all factors, we applied a quantile transformation to each spatial projections, replacing values by quantile ranks. This standardization places all outputs on a common scale, allowing us to identify both convergent diversity hotspots and regions of divergence across all above-mentioned factors. We refer to this standardized index as diversity index.

### 3.6. Post-processing of diversity estimates

#### 3.6.1. Global plankton diversity

To estimate global plankton diversity, we averaged all 10,080 projections generated by CEPHALOPOD, producing a mean diversity index equally weighted across observation types, Hill scaling factors, months, and bootstrap replicates. To propagate algorithm uncertainty into the global diversity estimate, we computed the standard deviation across bootstrap replicates for each grid cell, and averaged these values across observation types, Hill scaling factors and months. To assess the influence of observation types on diversity estimates, we calculated the standard deviation across observation types for each geographical cell, then averaged these values across Hill scaling factors, months, and bootstraps. Significance was tested per geographical cell, performing an ANOVA across values associated to each observation types. To assess how community evenness weighing influences global diversity indices, we first computed, for each Hill scaling factor (q), the mean diversity index averaged across observation types, months, and bootstrap replicates. Then we quantified the effect of increasing q by 1 in the estimated diversity index using a moving window of Δq = 1. Significance was tested per geographical cell, performing an ANOVA across values associated to each Hill scaling factor. To assess the distribution of diversity across the month dimension of our model outputs, we computed the difference between summer (*June, July, August*) and winter (*December, January, February*) months for each grid cell, then averaged these values across observation types, Hill scaling factors and bootstraps. Significance was tested per grid cell, performing an ANOVA between summer and winter values.

#### 3.6.2. Environmental drivers of diversity

To assess the environmental features explaining the variance in diversity in the trained algorithms, we averaged the variance explained by each environmental features across Hill scaling factors, months, and bootstrap replicates. We kept observation types separate in the variance explained as they relied on distinct modeling strategies: *predict first - assemble later* for occurrences and *assemble first - predict later* for abundance, biomass and metagenomic reads. This, and the fact that some features were not selected by CEPHALOPOD due to multicollinearity at the sampling locations (**Figure S8**, i.e., different across observation types) hinders intercomparability of the variance explained at the algorithm level. Therefore, we examine how environmental conditions structure the distribution of diversity indices across data archetypes by performing a Redundancy Analysis (RDA) at the projection level, including all diversity estimates and all environmental features tested. To assess how environmental conditions explain the diversity index change across in situ observation types and diversity metrics, we added both as supplementary variables to the RDA space. To ensure normality, the diversity indices projections were standardized by rescaling all values between 1 and 100 based on their rank (i.e., quantile transformation). From this RDA, we extracted the three first axes (RDA1, RDA2, RDA3) explaining 62, 7.5% and 3.2% of the variance, respectively, and the association strength of all environmental features to these axes (**Figure 3A**, **S7**).

#### 3.6.3. Biodiversity to ecosystem properties relationships

To quantify the strength of relationships between plankton diversity estimates and ecosystem properties, we performed pairwise Spearman correlations between the projected diversity indices, averaged by *in situ* observation types (i.e., the main axis of variance outside the LDG; see *section 2.3*.), and the ecosystem properties described in *section 2.1.5*. Then we quantified the relationship between diversity indices and each ecosystem property across observation types, by fitting a 2^nd^ degree polynomial regression that captures the response of tested ecosystem properties to diversity indices. The goodness of fit and uncertainty were estimated by computing an R^2^ and the associated standard deviation of the polynomial regression, respectively.

## STATEMENTS

## Acknowledgements

This project has received funding from the European Union’s Horizon 2020 Research and Innovation Programme under grant agreement no. 101094227 (Bluecloud2026) and no. 101059915 (BIOcean5D). This output reflects only the author’s view, and the European Union cannot be held responsible for any use that may be made of the information contained therein. The authors want to thank all the people involved in the Ocean Microbiomics and AtlantECO database initiative and related scientific cruises for making their data publicly available.

## Authors contribution

A.S. and M.V. conceived the study. M.V. supervised the study and provided fundings. A.S., F.B. and C.C. processed the input data. A.S. performed the analysis, wrote the first draft, and conceived the modelling pipeline. All authors substantially contributed to the successive versions of this manuscript.

## Competing interests

The authors declare there are no conflicts of interest for this manuscript.

## Funding source

Horizon 2020 Research and Innovation Programme, grant grant agreement no. 101094227 (Bluecloud2026) and grant agreement no. 101059915 (BIOcean5D).

## Data and code availability

All R scripts used for data preparation and model analysis are available at https://github.com/alexschickele/Diversity. All model outputs are available at https://doi.org/10.17882/112686. At time of publication, both will be publically available on Zenodo, under the Creative Commons Attribution 4.0 International license.

## Notes

https://www.seanoe.org/data/01015/112686/

### Competing Interest Statement

The authors have declared no competing interest.

