## Supplementary material for "Reconciling multiple views of global marine plankton diversity": Figure S

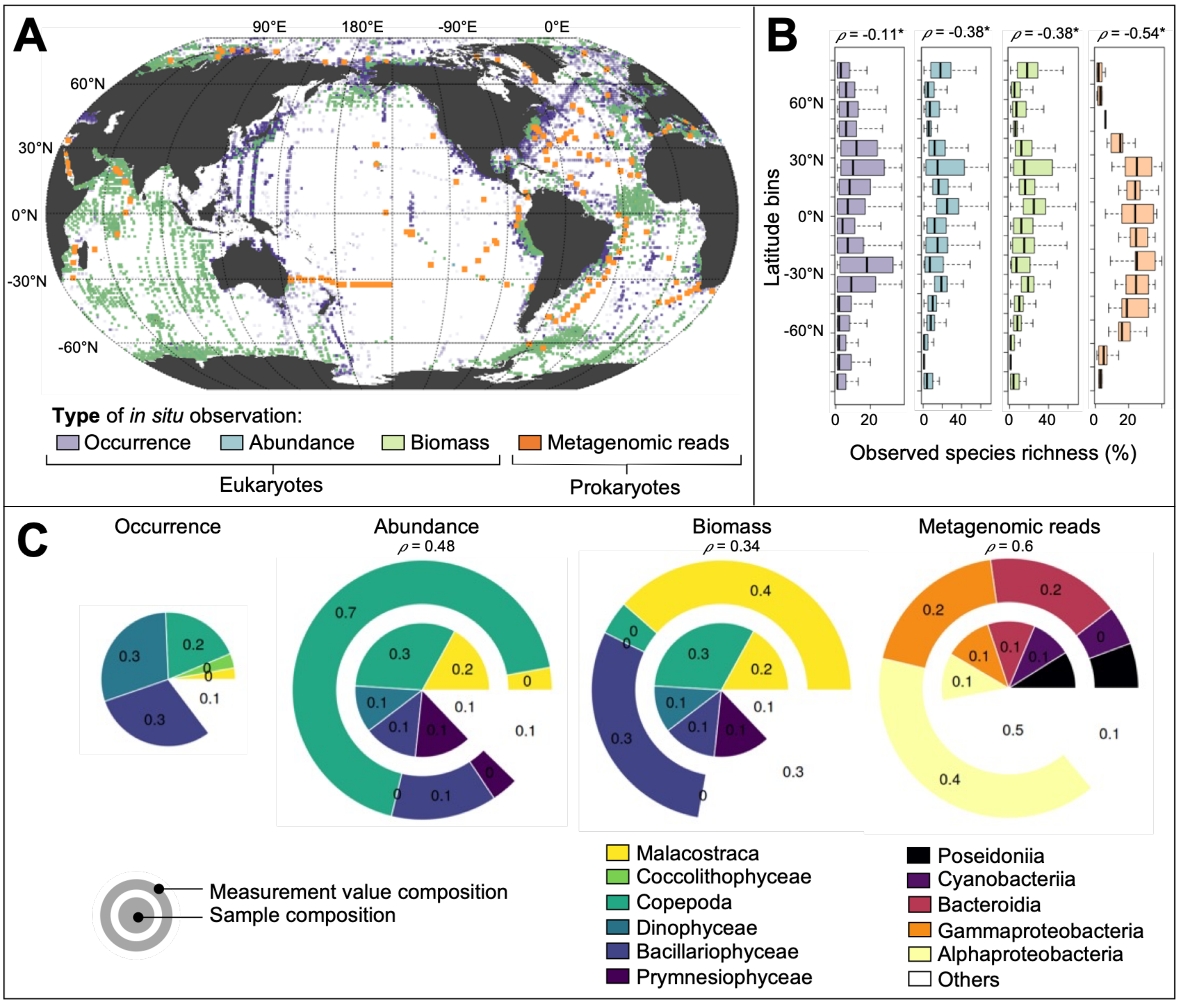


**Figure S1:** Plankton observation data retrieved from Vogt et al. (2026) and Paoli et al. (2022) with (**A**) sample location and (**B**) observed species richness pattern within 10° latitudinal bins, for Occurrence, Abundance, Biomass and Metagenomic are displayed together as species richness is independent of abundance or biomass measurements, that are associated with the same samples. The Spearman correlation between observed species richness and absolute latitudes is displayed above the boxplots (*p-value < 0.05). In panel (**C**) the inner pie chart shows the taxonomic composition in terms of number of species, while the outer pie chart represents the taxonomic composition weighted by the abundance, biomass or metagenomic reads (i.e., relative abundance of genes) associated with each species, aggregated at the class level. The Spearman correlation between the inner and outer pie charts is displayed above (*p-value > 0.05). Note that no outer pie chart is displayed for occurrence, as samples only contain presences (i.e., value of 1).

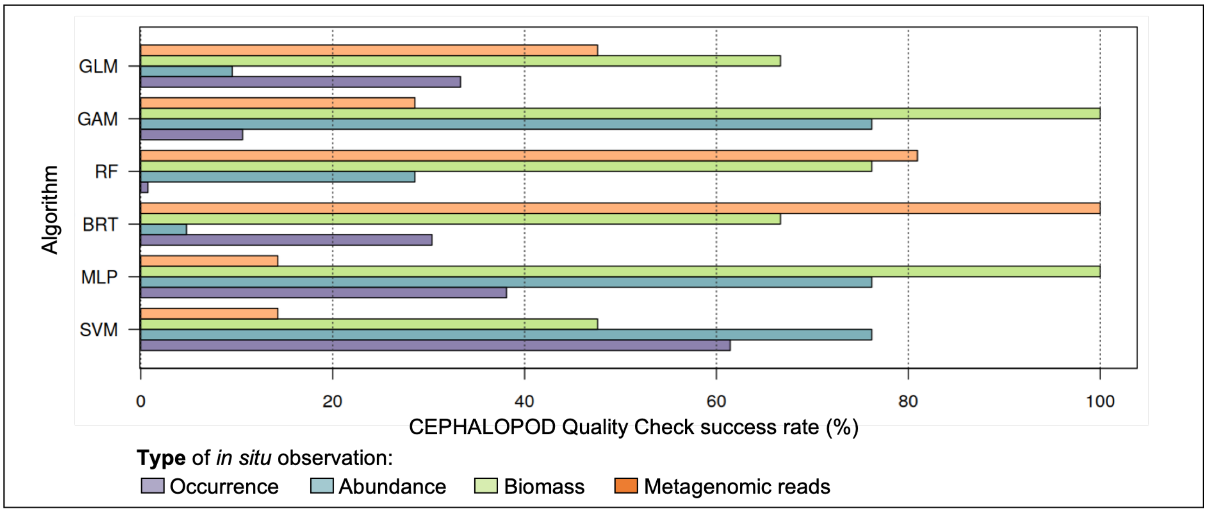


**Figure S2**: Quality check success for each algorithm trained in the CEPHALOPOD habitat modelling framework, averaged across all in situ observation types. The algorithm trained are Generalized Linear Model (GLM), Generalized Additive Model (GAM), Random Forest (RF), Booster Regression Tree (BRT), Multilayer Perceptron (MLP) and Support Vector Machine (SVM). The quality checks passed are (1) a priori variable importance, (2) predictive performance, (3) cumulative variance explained and (4) projection uncertainty. We refer to the method *section 3.4.* for details.

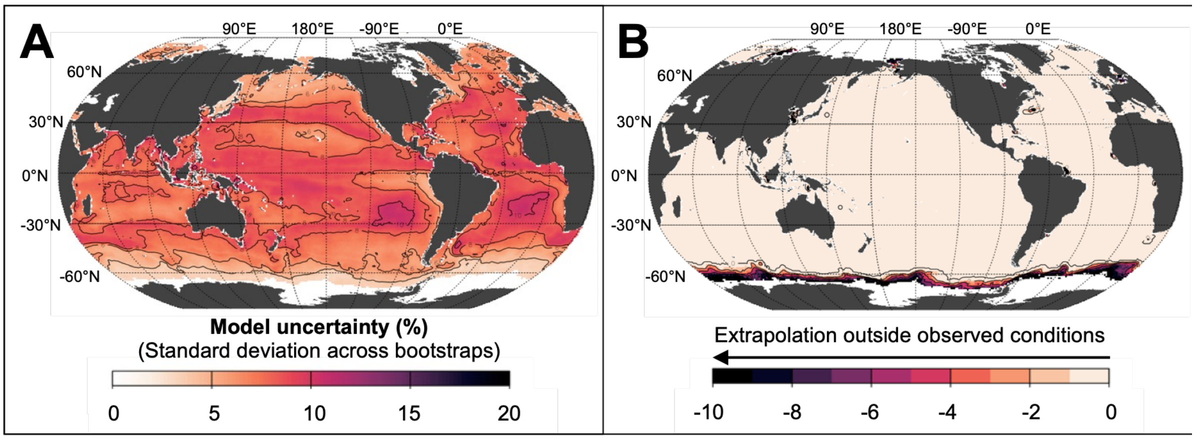


**Figure S3:** Global diversity index uncertainties with (**A**) the projected model uncertainty expressed as the standard deviation across bootstrap replicates normalized by the mean diversity index (%) and (**B**) the Multivariate Environmental Similarity Surface analysis, corresponding to model extrapolation outside observed conditions.

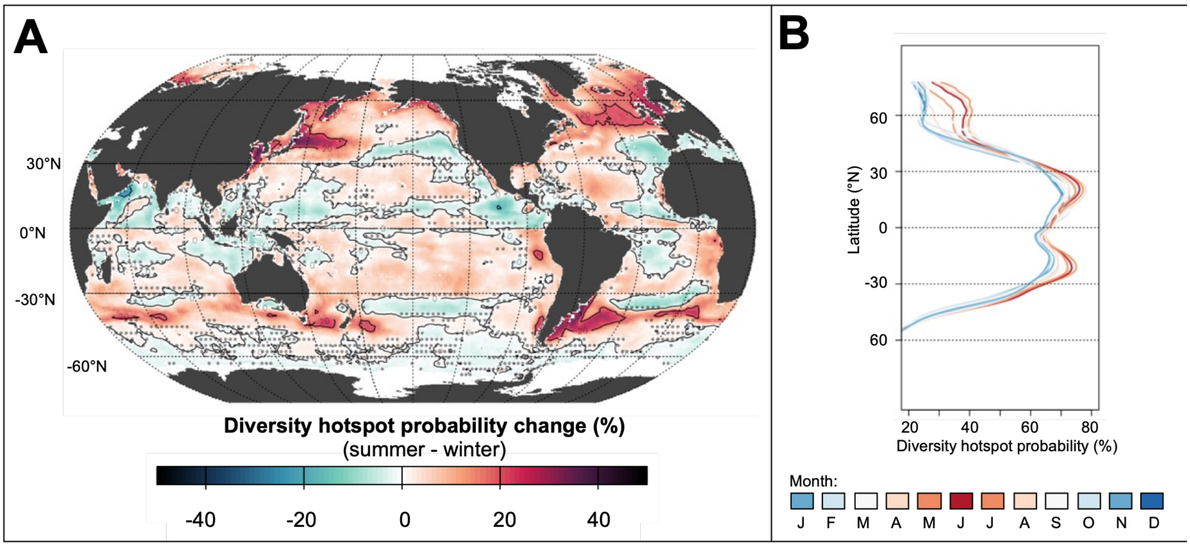


**Figure S4:** Diversity index change in function of summer and winter, with the global ocean projection (**A**) and latitudinal profiles per months (**B**). Black dots indicating no significant change, computed from an ANOVA across all values associated with a geographical cell (see method *section 3.6.1.*).

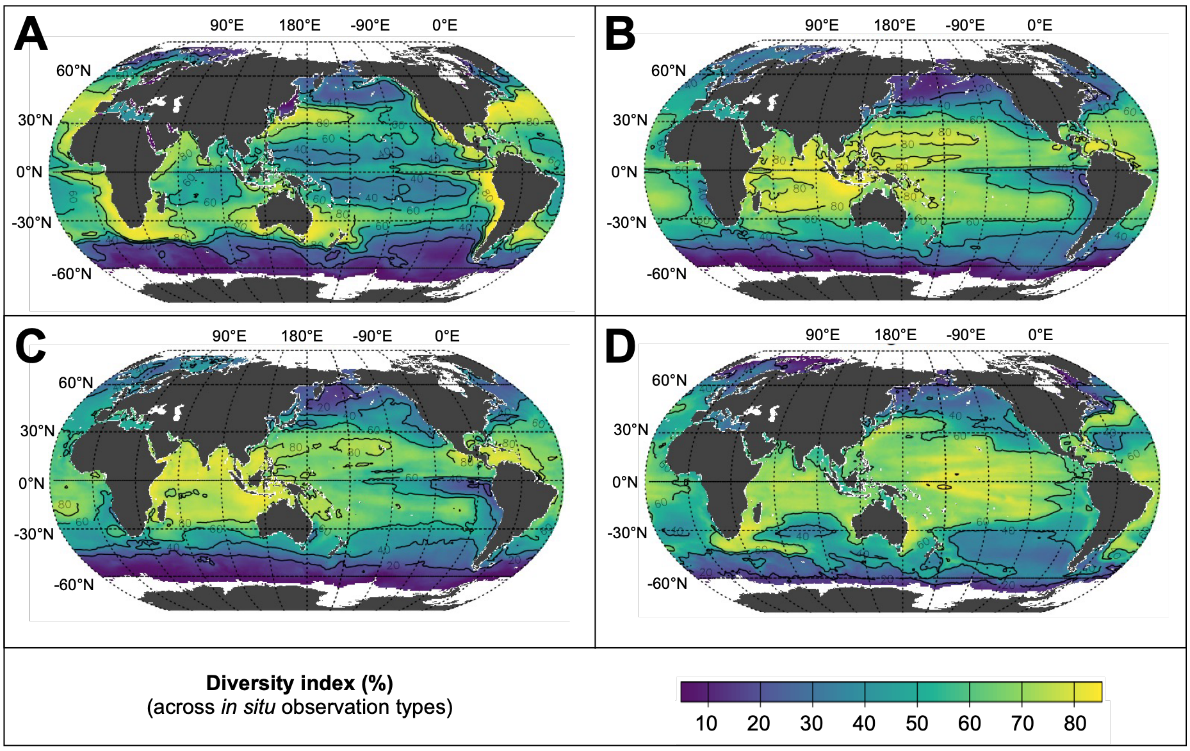


**Figure S5**: Diversity index averaged by in situ observation types, for (**A**) Occurrence, (**B**) Abundance, (**C**) Biomass and (**D**) Metagenomic reads data.

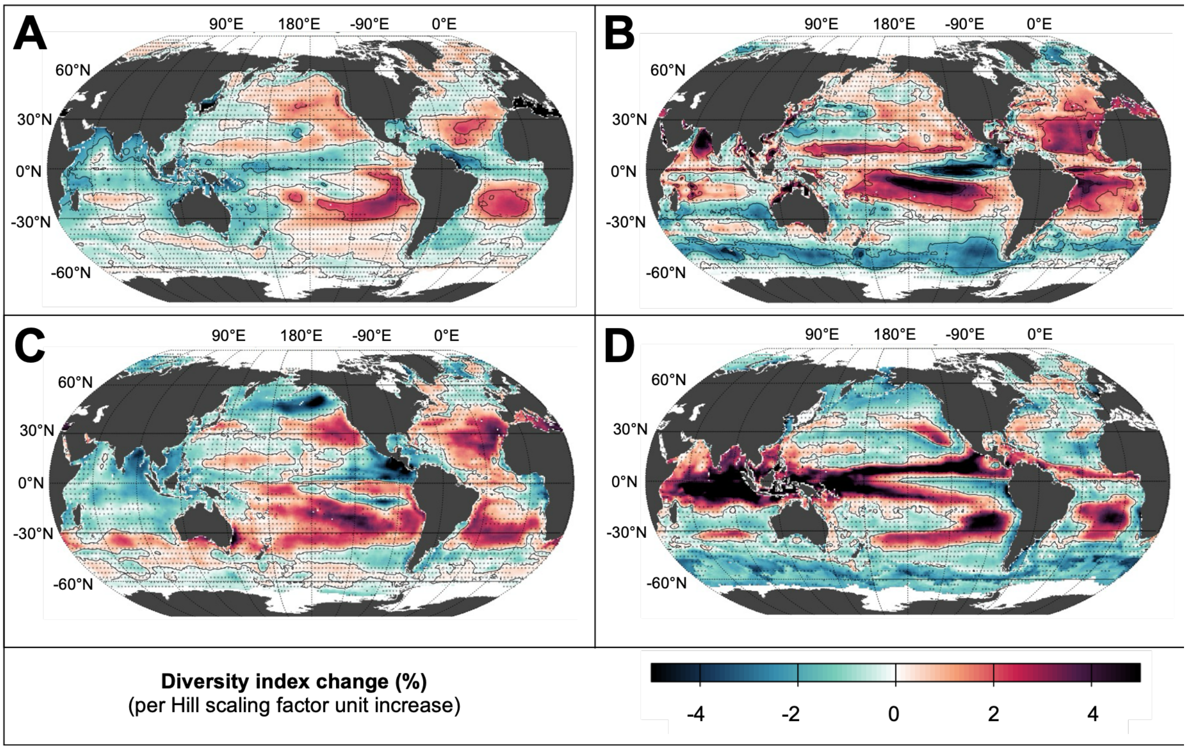


**Figure S6:** Diversity index change in function of Hill scaling factor, for (**A**) Occurrence, (**B**) Abundance, (**C**) Biomass and (**D**) Metagenomic reads data. Black dots indicating no significant change, computed from an ANOVA across all values associated with a geographical cell (see method *section 3.6.1.*).

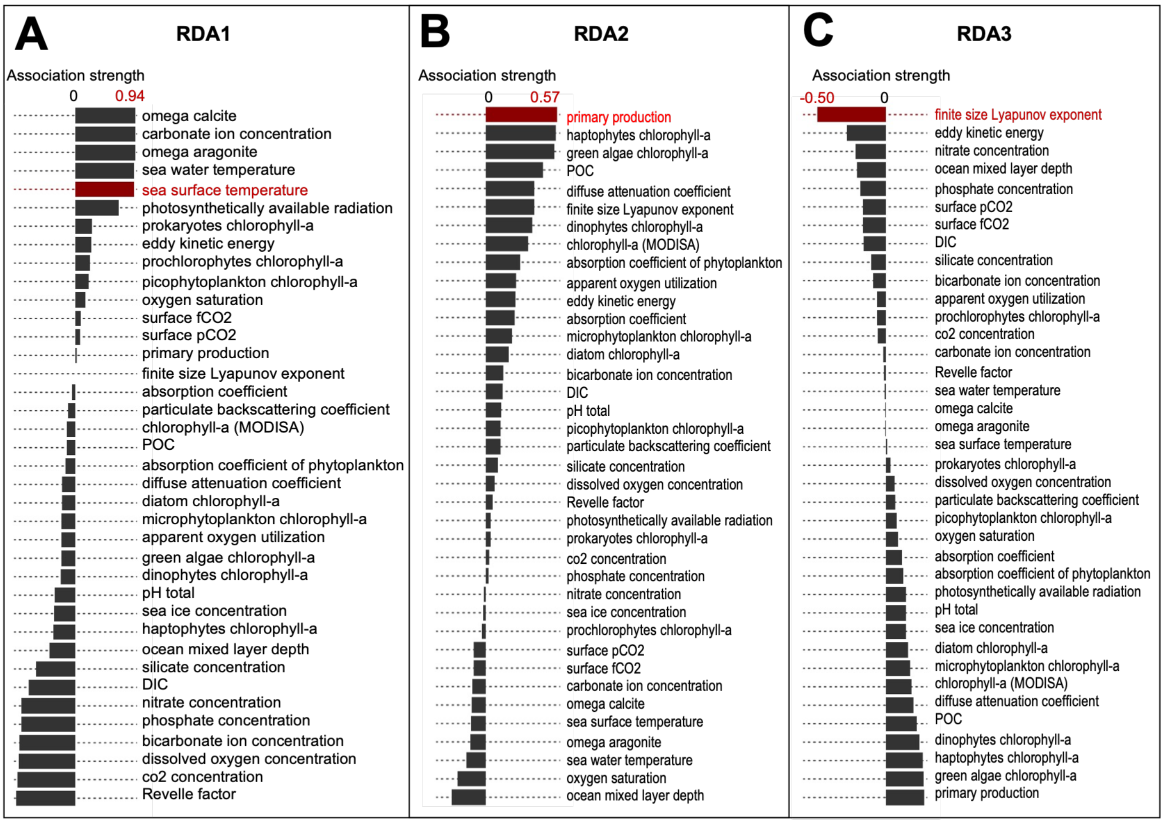


**Figure S7**: Redundancy Analysis (RDA) between projected diversity indices across *in situ* observation types, and hill scaling factors against environmental features. The environmental features associated to each of the two principal axes (RDA1, RDA2) are displayed as barplots, with the red bar indicating the environmental features considered as proxy of the RDA axes (i.e., temperature and primary production, respectively). The type of in situ observation is indicated by the shape of the points. The hill scaling factor is indicated by the color. For readability, only short labels are displayed for environmental features in the RDA space. We refer to **Table S1** for the correspondence between long and short names of environmental features.

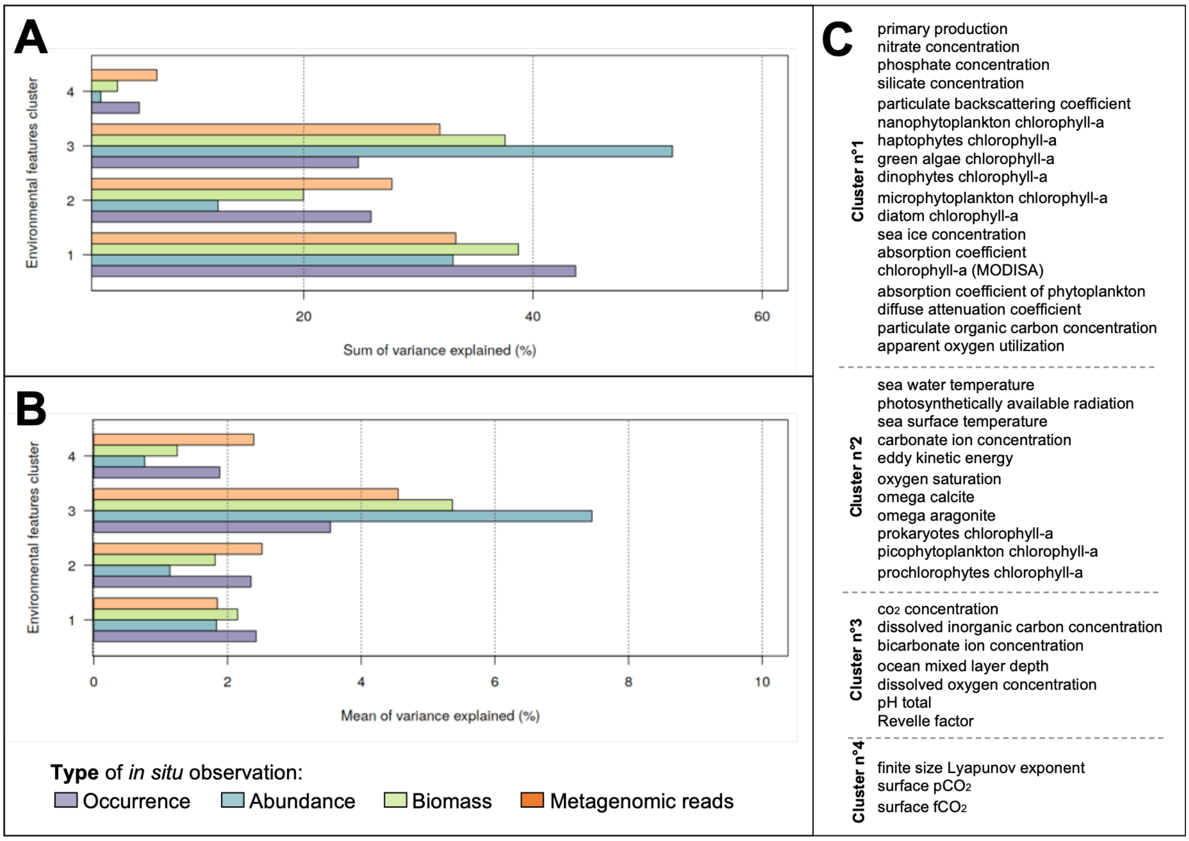


**Figure S8**: Variance explained (%) by each cluster of environmental features at the algorithm level, and partition across in situ observation types. Panel (**A**) and (**B**) indicate the sum and mean variance explained by environmental feature cluster, respectively. Note that the variance explained sums at 100% within each algorithm trained. Panel (**C**) indicates the environmental features associated to each cluster.

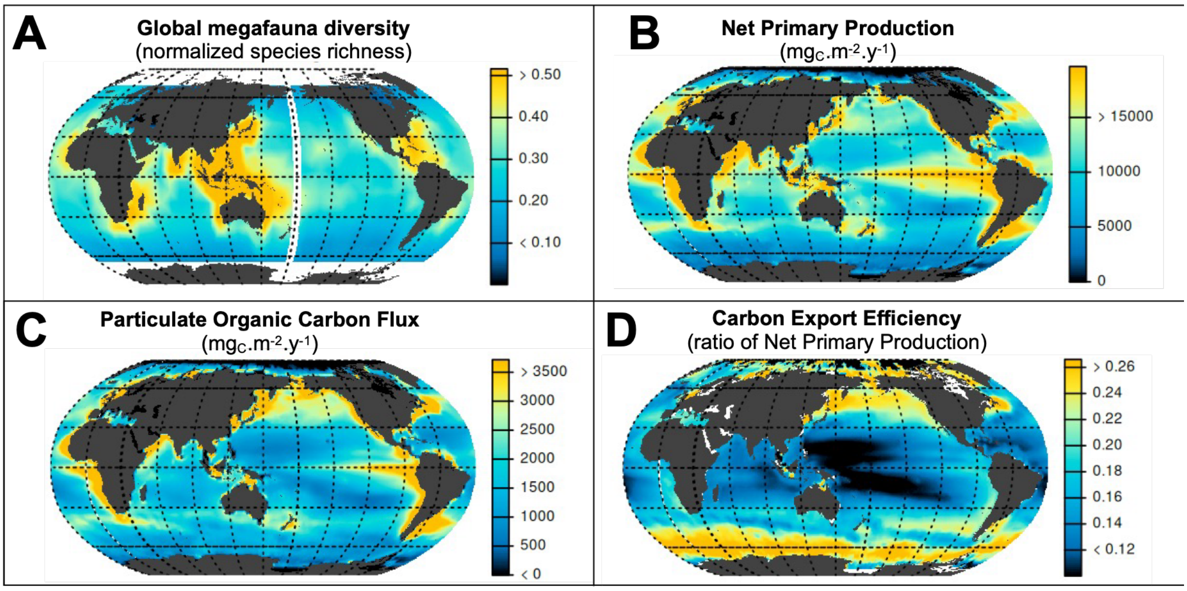


**Figure S9**: Global considered ecosystem properties with (**A**) Global oceanic megafauna biodiversity expressed as normalized species richness, (**B**) annual Net Primary Production (mgC.m$^{-2}$.d$^{-1}$), (**C**) the corresponding flux of particulate organic carbon (POC) exported below the euphotic zone (mgC.m$^{-2}$.d$^{-1}$) and (**D**) the corresponding efficiency of the production exported (POC/NPP; - ). Ecosystem properties are consider as presented in their reference papers, without spatial interpolation (Tittensor et al. 2010; DeVries and Weber, 2017).

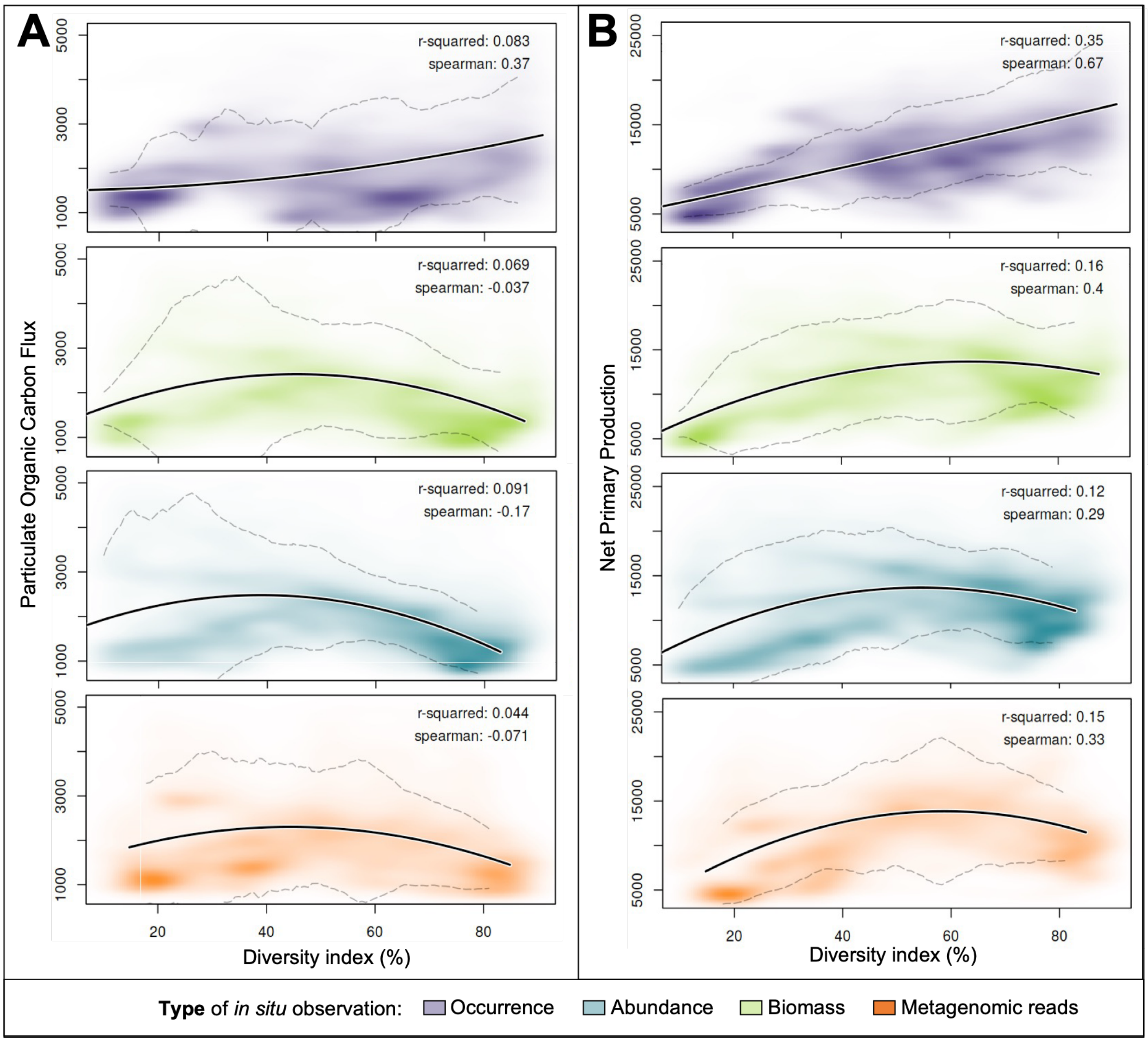


**Figure S10**: Emergent relationships between diversity index (%) and ecosystem functions for each in situ observation types, with (**A**) global flux of particulate organic carbon (POC) exported below the euphotic zone ($mgC.m^{-2}.d^{-1}$) and (**B**) annual Net Primary Production ($mgC.m^{-2}.d^{-1}$). The strength of the shading represents the density of geographical cells associated to a diversity index and ecosystem property bin in the global ocean. The continuous and dashed lines correspond to the average ± 1 standard deviation of a second order polynomial regression of the diversity index against the ecosystem property. The $R^2$ represent the goodness of fit of the regression while the spearman correlation indicates the strength of the linear trend.

**Table S1:** Collection of observation-based, monthly-resolved environmental features at the global scale derived from WOA18 (Garcia et al. 2018), Ocean Soda (Gregor and Gruber, 2020), GMIS (NASA, 2024), CMEMS (Xi et al. 2021) and AVISO (D'Ovidio, 2022).

| Long name (short name) | Unit | Source | Time range |
|---|---|---|---|
| ocean mixed layer thickness (M) | m | WOA18 | 1981-2010 |
| apparent oxygen utilization (A) | $\mu mol.kg^{-1}$ | WOA18 | 1900-2017 |
| silicate concentration (i) | $\mu mol.kg^{-1}$ | WOA18 | 1900-2017 |
| nitrate concentration (n) | $\mu mol.kg^{-1}$ | WOA18 | 1900-2017 |
| dissolved oxygen concentration (o) | $\mu mol.kg^{-1}$ | WOA18 | 1900-2017 |
| oxygen saturation (O) | % | WOA18 | 1900-2017 |
| phosphate concentration (p) | $\mu mol.kg^{-1}$ | WOA18 | 1900-2017 |
| sea water salinity (s) | - | WOA18 | 1955-2017 |
| sea water temperature (t) | °C | WOA18 | 1955-2017 |
| co2 concentration (co2) | $\mu mol.kg^{-1}$ | OceanSoda | 1982-2022 |
| carbonate ion concentration (co3) | $\mu mol.kg^{-1}$ | OceanSoda | 1982-2022 |
| dissolved inorganic carbon concentration (dic) | $\mu mol.kg^{-1}$ | OceanSoda | 1982-2022 |
| bicarbonate ion concentration (hco3) | $\mu mol.kg^{-1}$ | OceanSoda | 1982-2022 |
| sea ice concentration (ice) | - | OceanSoda | 1982-2022 |
| omega aragonite (omega_ar) | - | OceanSoda | 1982-2022 |
| omega calcite (omega_ca) | - | OceanSoda | 1982-2022 |
| pH total (ph_total) | -log([H+]) | OceanSoda | 1982-2022 |
| revelle factor (revelle_factor) | - | OceanSoda | 1982-2022 |
| surface fCO2 (fco2) | - | OceanSoda | 1982-2022 |
| surface pCO2 (pco2) | µatm | OceanSoda | 1982-2022 |
| absorption coefficient (443nm, QAA) (A_APD) | $m^{-1}$ | GMIS | 2002-2017 |
| absorption coefficient of phytoplankton (443nm, QAA) (A_APH) | $m^{-1}$ | GMIS | 2002-2017 |
| particulate backscattering coefficient (443nm, QAA) (A_ADC) | $m^{-1}$ | GMIS | 2002-2017 |
| chlorophyll-a (MODISA) (A_CHLA) | $mg\ Chl.m^{-3}$ | GMIS | 2002-2017 |
| diffuse attenuation coefficient (490nm, KD2) (A_K490) | $m^{-1}$ | GMIS | 2002-2017 |
| photosynthetically available radiation (400-700nm) (A_PAR) | $Ein\ m^{-2}.d^{-1}$ | GMIS | 2002-2017 |
| primary production (S_PP) | $mgC\ m^{-2}.d^{-1}$ | GMIS | 1997-2010 |
| particulate organic carbon concentration (POC) | $mgC.m^{-3}$ | CMEMS | 1998-2010 |
| diatom chlorophyll-a (DIATO) | $mgChl.m^{-3}$ | CMEMS | 1998-2020 |
| dinophytes chlorophyll-a (DINO) | $mgChl.m^{-3}$ | CMEMS | 1998-2020 |
| picophytoplankton chlorophyll-a (PICO) | $mgChl.m^{-3}$ | CMEMS | 1998-2020 |
| prochlorophytes chlorophyll-a (PROCHLO) | $mgChl.m^{-3}$ | CMEMS | 1998-2020 |
| prokaryotes chlorophyll-a (PROKAR) | $mgChl.m^{-3}$ | CMEMS | 1998-2020 |
| green algae chlorophyll-a (GREEN) | $mgChl.m^{-3}$ | CMEMS | 1998-2020 |
| microphytoplankton chlorophyll-a (MICRO) | $mgChl.m^{-3}$ | CMEMS | 1998-2020 |
| nanophytoplankton chlorophyll-a (NANO) | $mgChl.m^{-3}$ | CMEMS | 1998-2020 |
| haptophytes chlorophyll-a (HAPTO) | $mgChl.m^{-3}$ | CMEMS | 1998-2020 |
| eddy kinetic energy (eke) | $cm^2.s^{-2}$ | AVISO | 1993-2021 |
| finite size Lyapunov exponent (fsle) | $d^{-1}$ | AVISO | 2000-2021 |
